# Functional divergence of WWC family proteins in human endothelial cells

**DOI:** 10.64898/2026.08.21.746351

**Authors:** Tuli Pramanik, Austin Mills, Ondine Cleaver

**Affiliations:** Department of Molecular Biology, University of Texas Southwestern Medical Center, 5323 Harry Hines Blvd., Dallas, Texas, USA 75390

**Keywords:** WWC1, WWC2, WWC3, endothelial cell, Hippo, EMT, VEGFR2

## Abstract

The Hippo signaling pathway is increasingly recognized as a key regulator of endothelial cell (EC) proliferation, migration and vascular development. However, the roles of its upstream scaffold proteins remain poorly understood. Although WWC family proteins are widely regarded as functionally redundant activators of LATS1/2 kinases, the human genome contains a third family member, WWC3, that is absent from mice, raising the possibility of species-specific regulation of endothelial Hippo signaling. Here, we assessed the roles of WWC2 and WWC3 in human ECs using siRNA-mediated knockdown. Surprisingly, we found that WWC3 is the predominant regulator of canonical Hippo signaling, with a substantially greater effect than WWC2 on LATS1/2 phosphorylation, YAP/TAZ localization and expression of Hippo target genes. Loss of WWC3 also altered endothelial morphology and induced a partial endothelial-to-mesenchymal transition-like (EndoMT-like) phenotype. By contrast, WWC2 had a lesser effect on canonical Hippo signaling, but it was required for normal VEGF signaling dynamics. Despite these distinct molecular functions, depletion of either WWC2 or WWC3 impaired EC proliferation, migration, and cord formation in vitro. Together, our findings demonstrate that WWC family proteins perform overlapping but distinct functions in human ECs, with WWC3 acting as the predominant canonical Hippo regulator, whereas WWC2 more efficiently modulates VEGF signaling. These results reveal unexpected functional specialization among WWC proteins and suggest that regulation of Hippo signaling in human ECs differs from that inferred from mouse studies.

**HIGHLIGHTS:**

- WWC2 and WWC3 are expressed in endothelial cells.
- WWC3 acts as a canonical facilitator of LATS1/2 phosphorylation and YAP/TAZ localization.
- WWC2 and WWC3 both regulate migration, tube formation, and proliferation.
- WWC2 regulates the duration of VEGF signaling in human ECs.
- WWC3 loss-of-function causes endothelial cells to upregulate mesenchymal genes.

## INTRODUCTION

Endothelial cells (ECs) continuously integrate biomechanical, biochemical and cell-cell junctional signals to regulate their proliferation, migration, morphology and barrier function.^1,2^ These coordinated responses are essential for angiogenesis during development and for vascular homeostasis throughout life. Among the signaling pathways that govern endothelial behavior, the Hippo pathway has emerged as a central regulator of endothelial growth, mechanotransduction and vascular remodeling.^3,4^ Studies of the core Hippo kinase cascade, including LATS1/2 and the transcriptional coactivators YAP1 and TAZ, have established critical roles for this pathway in endothelial specification, angiogenesis and vascular integrity.^5,6^ Despite these advances, considerably less is known about the upstream proteins that regulate Hippo pathway activity in ECs. Understanding how these upstream regulators integrate extracellular signaling into Hippo signaling is essential for defining the mechanisms that coordinate vascular growth, remodeling and homeostasis.

Genetic studies have established that the core Hippo components LATS1/2 and the downstream transcriptional coactivators YAP1 and TAZ are essential for angiogenesis, vascular remodeling and endothelial mechanotransduction. In the canonical pathway, LATS1/2 phosphorylate YAP1 and TAZ, promoting their cytoplasmic retention and limiting transcriptional activation of pro-growth target genes.^7,8^ However, accumulating evidence suggests that Hippo pathway components also perform endothelial functions that cannot be fully explained by regulation of YAP1/TAZ alone.^5,9,10^ For example, the upstream regulator Merlin (NF2) suppresses excessive vascular sprouting by modulating VEGFR2 trafficking independently of YAP1/TAZ, while LATS1/2 regulate endothelial responses to biomechanical forces through both YAP1-dependent and YAP-independent mechanisms.^9^ Taken together, these studies suggest that upstream regulators of the Hippo pathway may have specialized functions in ECs beyond simply controlling YAP1/TAZ activity. However, the identity and endothelial-specific roles of these upstream regulators remain poorly understood.

Among the best characterized upstream regulators of Hippo signaling are the WWC family of scaffold proteins (WWC1/KIBRA, WWC2 and WWC3), known to promote the activation of LATS1/2 via direct protein-protein interactions.^11–14^ Biochemical studies have shown that each WWC family member is capable of activating LATS kinases, leading to the prevailing view that these proteins function largely redundantly. In vivo, however, mouse studies have identified an essential role for WWC2 during embryonic vascular development and postnatal angiogenesis, whereas loss of WWC1 does not lead to overt vascular defects.^15,16^ Whether this reflects functional specialization among WWC family members or simply differences in their expression remains unknown. Importantly, unlike in mouse, humans have a third WWC family member, WWC3.^17^ This raises the possibility that Hippo pathway regulation in human ECs differs from that inferred from mouse studies.

Here, we investigated the expression and function of WWC family proteins in human ECs. We found that WWC2 and WWC3 are both expressed in human ECs but perform distinct functions. We show that WWC3 is the predominant regulator of canonical Hippo signaling, controlling LATS1/2 phosphorylation, YAP1/TAZ activity, and maintenance of endothelial identity. By contrast, WWC2 has relatively modest effects on canonical Hippo signaling but regulates VEGF signaling dynamics. Despite these distinct molecular functions, depletion of either WWC2 or WWC3 impairs fundamental EC behaviors, including proliferation, migration and cord formation. Taken together, our results demonstrate functional specialization among WWC family members and suggest that regulation of Hippo signaling in human ECs is more complex than previously appreciated from mouse studies.

## RESULTS

### Expression of the WWC family of proteins in endothelial cells

To understand the role of WWC family proteins in endothelial cells (ECs), we investigated expression and function of WWC1, WWC2 and WWC3 in Human Umbilical Vein Endothelial Cells (HUVECs). First, using the publicly available comprehensive Human Vascular Cell Atlas,^18^ we assessed transcripts for all 3 members of this family. We found that only *WWC2* and *WWC3* were expressed at detectable levels in human ECs (**Fig. 1A**). Examination of the distribution of both WWC2 and WWC3 revealed that they were relatively broadly expressed in adult human arterial, venous, capillary and lymphatic ECs, as well as mural cells, when assessing the database clustering information of pooled ECs from multiple ages, genders and organs (**Fig. 1A’**). Comparing with the expression of brain EC-enriched genes (*SLC2A1*, *MFSD1A* and *SLC7A5)*, or those enriched in liver (*CLEC4M*), lung ECs (*HPGD*) or spleen ECs (*CD8A*), we were able to identify organotypic capillary clusters (**Fig. 1B**). This analysis revealed that *WWC2* was particularly enriched in human brain capillaries, but lower in the vessels of other tissues. *WWC3*, by contrast, is broadly expressed throughout the endothelial tree.

**Figure 1.**
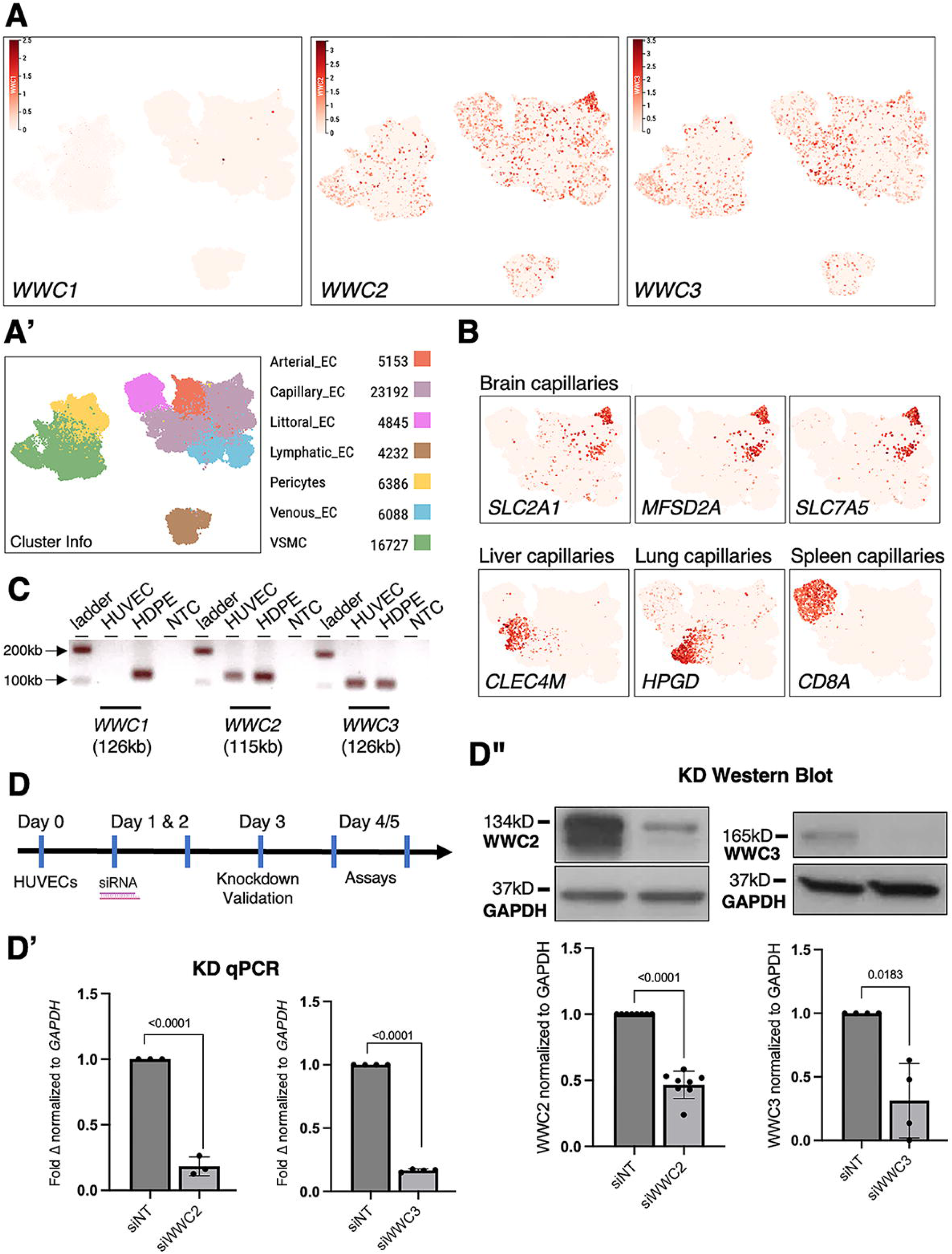
Endothelial expression and siRNA-mediated knockdown of WWC2 and WWC3. **A,A’)** *WWC1*, *2*, 3 expression per publicly available data on the Vascular Cell Atlas (https://www.vascularcellatlas.org). **B)** Tissue specificity of EC clusters shown using genes known to be expressed in brain capillaries (*SLC2A1*, *MFSD2A* and *SLC7A5*), or liver (*CLEC4M*), lung (*HPGD*) and spleen (*CD8A*) capillaries. **C)** RT-PCR shows *WWC2* and *WWC3* expression in HUVECs, whereas *WWC1* is not expressed highly in HUVECs. HDPE was used as a positive control for WWC expression (NTC: No template control). **D**) Schematic of siRNA-mediated knockdown system used to study the role of *WWC2* and *3* in ECs. **D’)** qPCR and **D’’)** Western Blot shows successful knockdown (KD) of *WWC2* and 3 in ECs. Significance was calculated using unpaired Student’s t-test.

To validate expression, RT-PCR was carried out on cDNAs generated from HUVECs and control HDPE cells. We found that *WWC2* and *WWC3* were both expressed in HUVECs and control Human Pancreatic Ductal Epithelial cells (HDPEs), while *WWC1* expression was only observed HDPEs (**Fig. 1C**), confirming data from the human EC Atlas. Similarly, *WWC2* and *WWC3*, but not *WWC1*, were also expressed in Human Pulmonary Arterial ECs (HPAECs), pancreatic islet ECs (Mile Sven 1, or MS1s), and Human Aortic ECs (HAECs) (data not shown).

To assess whether either *WWC2* or *WWC3* was required in HUVECs, we used an siRNA-mediated knockdown (KD) approach (**Fig. 1D**). We confirmed expression reduction, both at the transcript and protein level. First, we carried out qPCR on cDNA from cells treated with negative control siRNA (non-targeting, siNT) or *WWC2*- or *WWC3*-targeting siRNA (siWWC2 or siWWC3) to knockdown endogenous expression. We observed an approximately 82% reduction in *WWC2* mRNA levels and 83% reduction in *WWC3* mRNA levels (**Fig. 1D’**). These data were further validated using western blot analysis with antibodies to WWC2 and WWC3, following the same 3-day siRNA regimen (**Fig. 1D”**).

### WWC3 is the predominant canonical regulator of Hippo signaling in human ECs

Previous studies have shown that Wwc2 null embryos exhibit an increase in YAP1/TAZ target gene expression in whole embryo RNAseq,^15^ as well as marked vascular defects.^16^ We therefore analyzed expression of the known YAP1/TAZ target genes *CTGF*, *CYR61*, *ANKRD1*, and *TGFβ2* in siWWC2 and siWWC3 cells 48hrs after knockdown (**Fig. 2A**). As expected, upon loss of either *WWC2* or *WWC3* expression, we found a concomitant increase in expression of these genes (**Fig. 2B**). However, loss of WWC3 increased *TGF*β2 expression approximately 12-fold, which was dampened when both WWC2 and 3 were co-depleted. Thus, our results confirm WWC2 and WWC3 regulation of Hippo targets, validating previous findings. However, our studies in HUVECs suggest that WWC3 has a more significant impact on classical Hippo target than WWC2.

**Figure 2.**
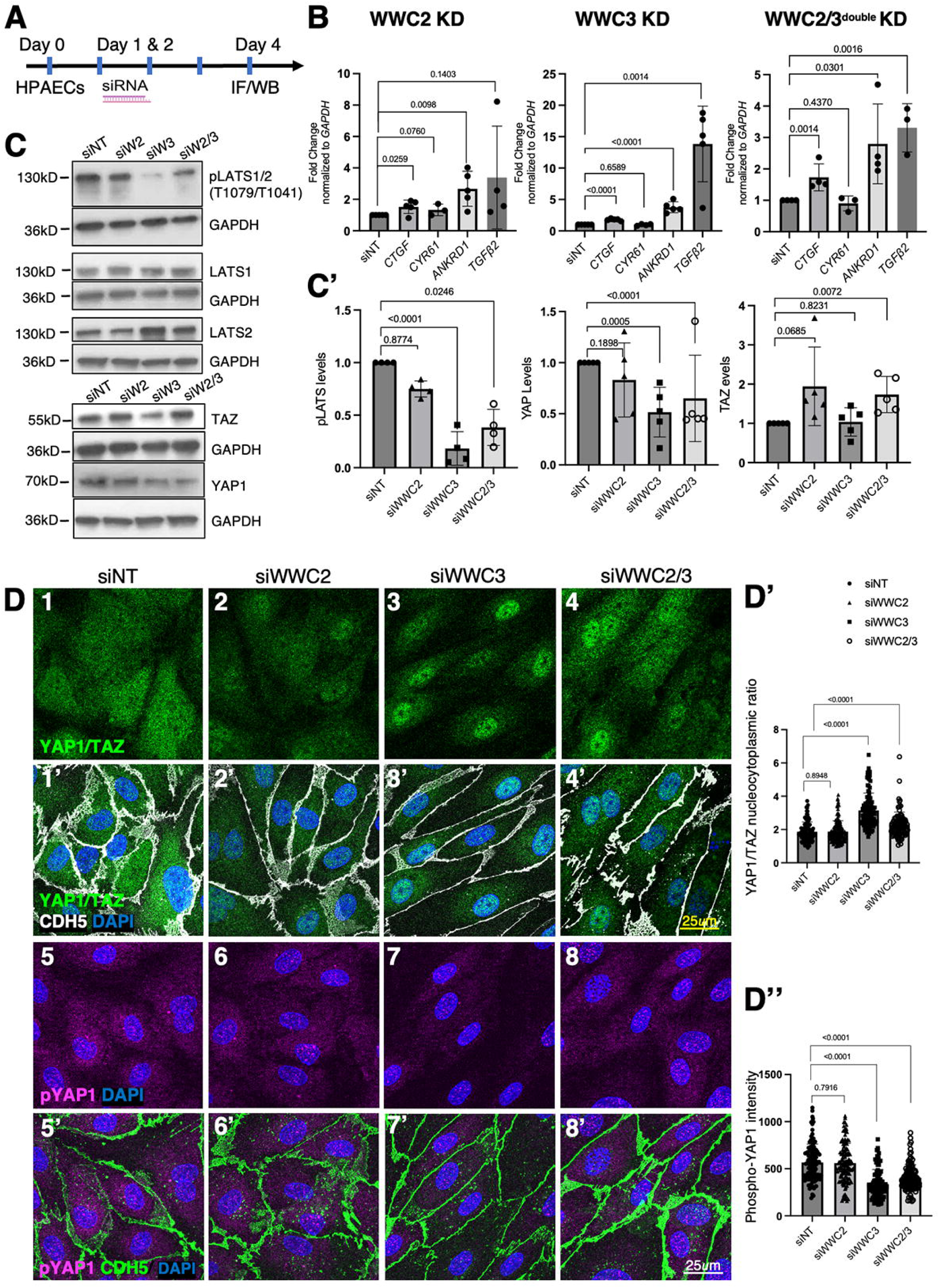
WWC3 is the predominant canonical regulator of Hippo signaling in ECs. **A)** Schematic for siRNA-mediated knockdown of *WWC2*, *WWC3* and *WWC2/3* in ECs. **B)** Loss of *WWC2* and *3* lead to increase in YAP1/TAZ target genes like *CTGF*, *ANKRD1*, and *TGFB2*, but not *CYR61.* **C)** Western Blot shows sharp reduction in pLATS in siWWC3, but lesser in siWWC2 cells. In addition, the effect is blunted in siWWC2 treated cells. Loss of WWC3 also impacts total levels of YAP1 and TAZ. siWWC2 and siWWC2/3 shows a blunted reduction in TAZ, but not YAP1. **C’)** Quantification of Western bands in C. **D)** siWWC3 and siWWC2/3 exhibit higher nuclear YAP1 and reduction in pYAP1. Panels 1-4 IF with antibody that recognizes both YAP1 and TAZ. Panels 1’-4’ same panels as 1-4 but with CDH5 IF. Panels 5-8 antibody stain for pYAP1. Panels 5’-8’ same panels as 5-8 but with CDH5 IF. These data are quantified in D’ and D”, respectively. Significance was calculated using unpaired Student’s t-test. Scale bars: All panels 25μm.

Given the known role of WWC proteins as scaffolds to facilitate LATS1/2 phosphorylation,^11,13,19^ and the finding that expression of any WWC proteins can upregulate LATS1/2 phosphorylation (pLATS1/2) in HEK293A cells,^12^ we tested the role of WWCs in HUVECs. Using siRNA approaches, we assessed the status of pLATS1/2 upon depletion of WWC2, WWC3 or WWC2/3 together. Surprisingly, we observed a significant reduction of LATS1/2 phosphorylation in siWWC3 cells (80%), but not siWWC2 (**Fig. 2C, C’**). Depletion of WWC2 only mildly reduced phosphorylation of LATS1/2 (25%), but it was not statistically significant. Loss of pLATS1/2 was dampened when both WWC2 and 3 were knocked down together (60%).

Interestingly, loss of WWC proteins impacted total levels of YAP1 and TAZ protein. Depletion of WWC3 led to an almost 50% decrease in the total levels of YAP1, but not TAZ; while depletion of WWC2 led to a significant increase in the total level of TAZ, though levels of YAP1 remained unchanged (**Fig. 2C, C’**). These findings are notable in that most studies discuss nucleocytoplasmic localization of YAP1 and TAZ, rather than protein levels.

We next tested whether the loss of LATS1/2 phosphorylation affected YAP1/TAZ localization in siWWC2 and siWWC3 cells. We used immunofluorescence analysis (IF) using antibodies that recognize either both YAP1 and TAZ (**Fig. 2D1**), or pYAP1 alone (**Fig. 2D3**), as well as junctional vascular endothelial cadherin 5 (VEcad or CDH5) to outline EC boundaries (**Fig. 2D2,4**). In control HUVECs (siNT-treated) plated at high confluency (where all cells touch adjacent cells), YAP1 and TAZ were localized in the cytoplasm (**Fig. 2D**), while the cells treated with siWWC3 showed an increase in nuclear-to-cytoplasmic ratio (70%), the double knockdown blunted this increase (20%). Interestingly, we did not observe any significant change in YAP/TAZ localization when cells were treated with siWWC2. The increase in YAP1/TAZ nuclear-to-cytoplasmic ratio thus correlated with reduced LATS1/2 phosphorylation. Nuclear YAP1/TAZ was markedly higher upon loss of WWC3 (60%) (**Fig. 2D’**). Along those lines, we also stained siRNA-treated cells with anti-phospho-Yap1 (pYap1) and observed significantly reduced levels of cytoplasmic pYap1 in siWWC3 and siWWC2/3 cells, demonstrating that WWC3 impacts activity of Yap1 more robustly (**Fig. 2D’’**).

Taken together, these results suggest that while WWC3 functions as the predominant canonical regulator of LATS1/2 phosphorylation, impacting cellular localization of YAP1/TAZ and increasing expression of their targets, WWC2 does so as well albeit less robustly. In addition, loss of WWC2 actually increases the levels of TAZ protein in ECs, with less of an effect on the nuclear-to-cytoplasmic ratio of YAP1/TAZ.

### WWC proteins are required for fundamental EC behaviors

To determine whether WWC2 and WWC3 are required for fundamental EC behaviors, we assessed their roles in proliferation, migration and cord formation in vitro. Because the Hippo pathway has been shown to impact cell proliferation,^20^ we first tested whether loss of WWC2 or WWC3 affected this process in ECs. Using IF staining for Ki67 and CDH5, we observed a modest but significant reduction in Ki67-positive nuclei in WWC2 knockdown ECs (20% reduction), while loss of WWC3 resulted in more robust reduction (90% reduction) (**Fig. 3A, A’)**. We also noted that loss of WWC3 led to dramatic alteration of EC cell shape, with ECs adopting a spindle-like shape.

**Figure 3.**
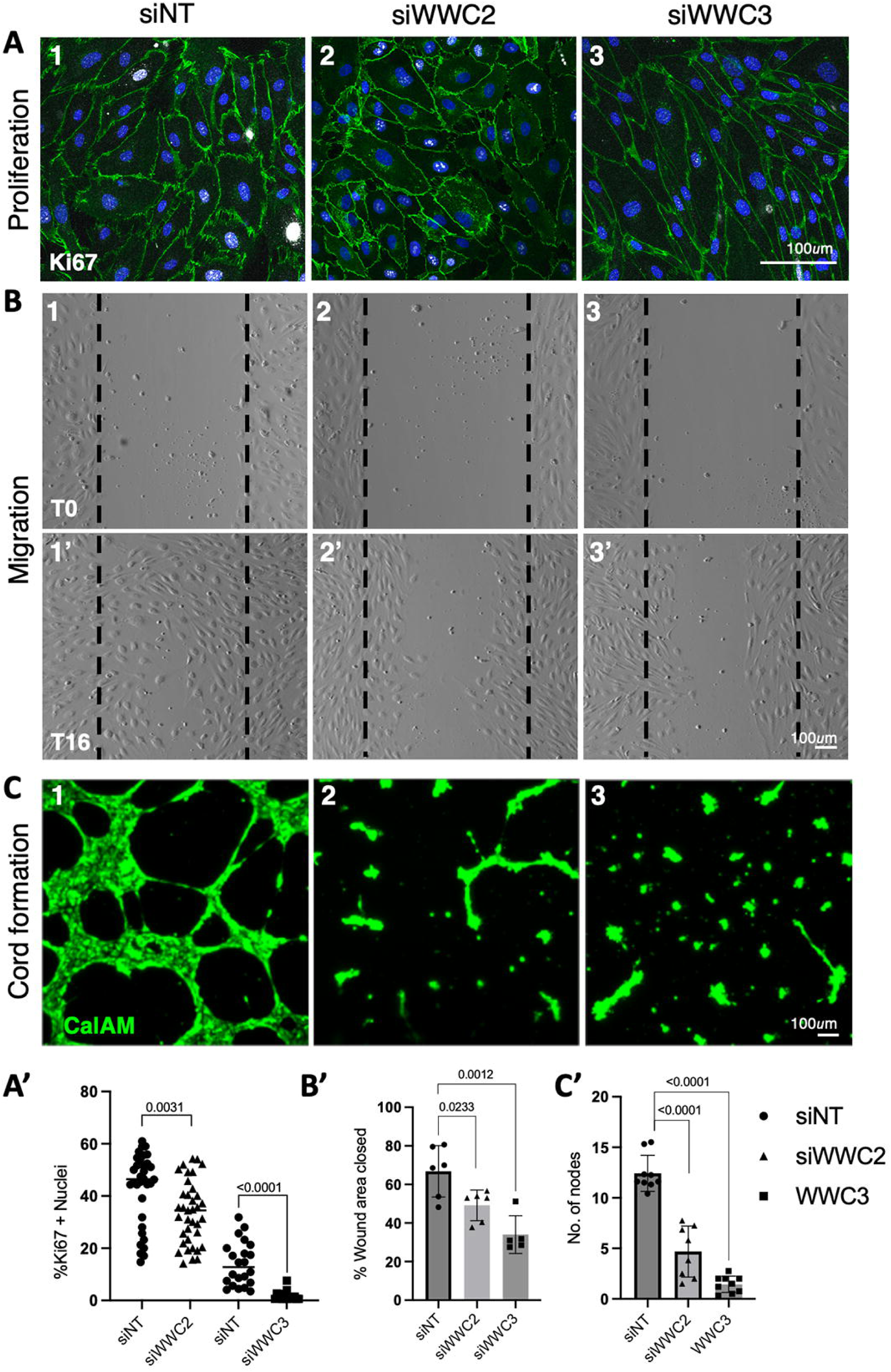
WWC2 and WWC3 are required for endothelial cell proliferation, angiogenic cord formation, and migration. **A)** Ki67 staining shows the reduction in proliferation in *WWC3* knocked down cells, quantified in A’. **B)** Wound healing assays show the failure of *WWC2* and *WWC3* knocked down cells to migrate and close the wound 16hrs post wound creation (T16hr timepoint shown in panels 1’-3’), quantified in **B’**. **C)** Angiogenesis assay shows the failure of ECs to form networks of cords in the absence of WWC2 and WWC3, quantified in **C’**. Significance was calculated using unpaired Student’s t-test. Scale bars: All panels 100μm.

We next tested the effect of WWC2 or WWC3 loss on the migratory ability of HUVECs using a standard *in vitro* wound healing assay.^21^ In this assay, ECs are plated to confluence, and a scratch or ‘wound’ is created within the center of the field of cells, allowing a clear area that ECs can migrate into. We found that loss of either WWC2 or WWC3 resulted in decreased migration of ECs, as ECs filled the empty area less efficiently than controls after 16hrs of culture (**Fig. 3B**). The percent area of migration/wound closure in both siWWC2 and siWWC3 cells decreased significantly when compared with siNT-treated cells (**Fig. 3B’**).

Lastly, we tested the role of WWC2 and WWC3 in classic ‘angiogenesis’ assays.^21^ One of the characteristics of ECs in culture is their ability to form a plexus of cords when plated on Matrigel, mimicking the in vivo phenomenon that precedes opening of lumens during early development (or cord formation). We carried out this angiogenesis assay on control and siRNA-treated cells to test the role of WWC proteins on this process. Following 12hrs of culture after plating, we observed a significant decrease in cord formation upon loss of either WWC2 or WWC3, with fewer connections between clusters of ECs, as shown by the live cell stain calcein acetoxymethyl ester (CalAM) (**Fig. 3C, C’**). Together, these three assays revealed the requirement for WWC2 and WWC3 in EC basic biological processes.

### WWC2 is essential for the duration of endothelial VEGF signaling in human ECs

Studies of the global *Wwc2^-/-^* reported severe vascular defects, which were accompanied by sharp elevation of VEGF-A, a growth factor that drives angiogenesis.^15^ However, this global, whole embryo *Wwc2* null could not distinguish the role of WWC2 specifically in ECs versus other cell types. Given that the Hippo pathway has been implicated in regulating VEGF-VEGFR2 signaling, we tested whether WWC family proteins might similarly influence this signaling axis.^22^

Using recombinant human VEGF-A and cultured HUVECs, we asked how endothelial loss of WWC2 or WWC3 affected VEGF-A signaling. As indicated in the experimental schematic (**Fig. 4A**), siWWC2 and siWWC3 treated cells were starved and then treated with 50ng/mL of VEGF at the indicated time points. VEGF signaling was assessed by changes in VEGF receptor 2 (VEGFR2) phosphorylation (pVEGFR2), as well as downstream signaling via phospho-AKT (pAKT) and phospho-ERK1/2 (pERK1/2). We found that while VEGF signaling in control siNT cells showed measurable phosphorylation of VEGFR2, AKT and ERK shortly after VEGFA addition, that signaling was attenuated after 30min and even more so at 60min, with pAKT being gone entirely at that timepoint (**Fig. 4B**). By contrast, loss of WWC2 led to an abnormally prolonged response, with significant signal at 30min and high levels of both pVEGFR2 and pERK1/2 at 60min. In contrast, loss of WWC3 did not produce significant elevation in signaling at 30min, although residual elevation of both pVEGFR2 and pERK1/2 continued to 60min. These data suggest a slightly more prominent role for WWC2 in VEGFA signaling.

**Figure 4.**
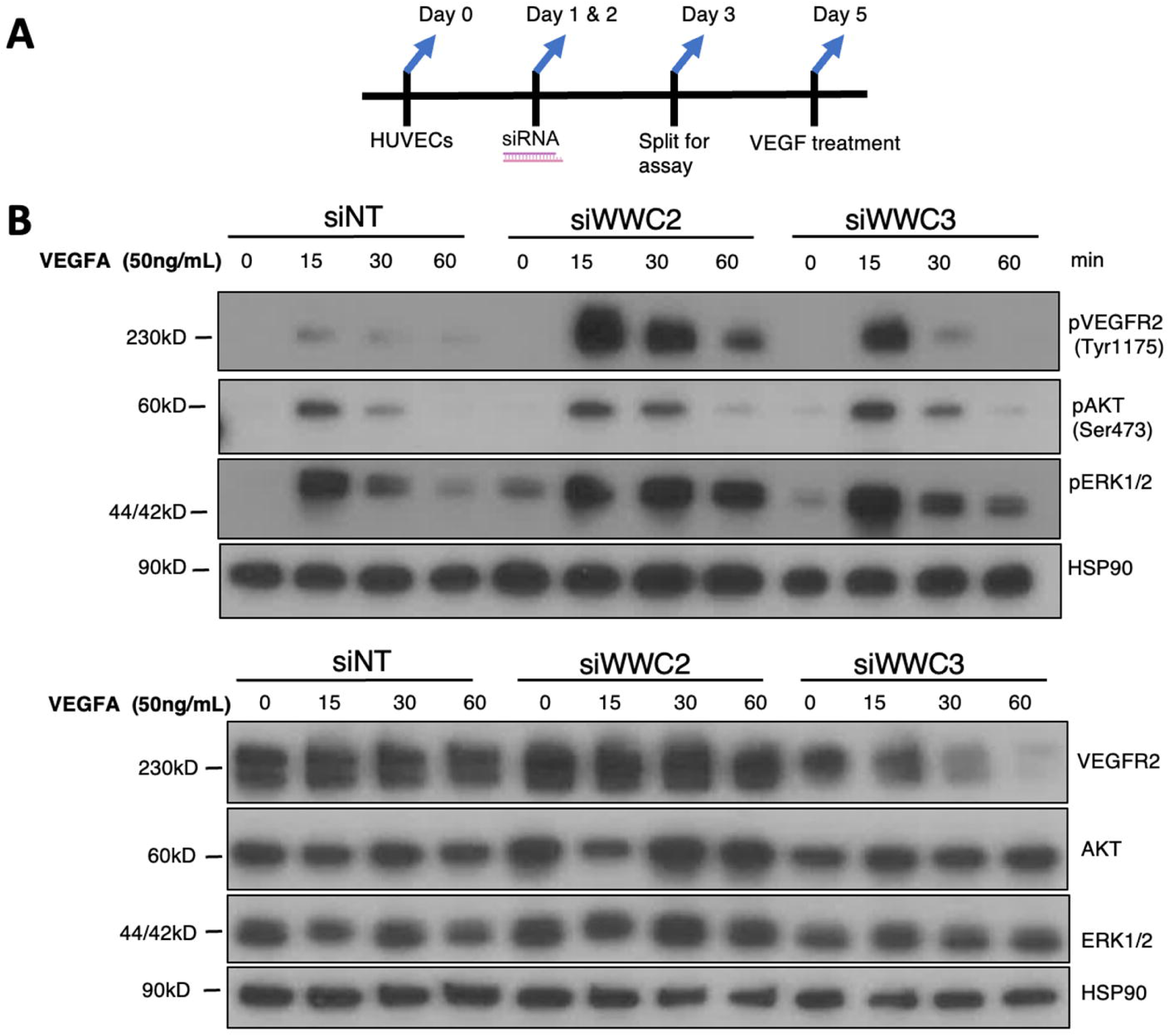
WWC2 is essential for proper endothelial VEGF signaling. **A)** Schematic showing how ECs were treated with siRNA, split and exposed to 50ng/mL of recombinant human VEGF, at indicated timepoints, to test VEGFR2 activation. **B)** Analysis of phospho-VEGFR2, and downstream activation of AKT and ERK pathways, was carried out by Western blotting to example response downstream of VEGF in the presence or absence of WWC2 and WWC3.

### Loss of WWC3, but not WWC2, impacts EC cell shape

Interestingly, we observed that loss WWC3 in ECs led to dramatic morphological changes, while the loss of WWC2 does not (**Figs. 2 and 3**). This was evident siWWC3-treated cells in brightfield (**Fig. 5A**) which showed elongation of cells and a spindle-like appearance in. In addition, despite confluency remaining equal between control and siRNA samples, we noted a distinct “swirling” organization of siWWC3-treated cells. CDH5 staining of control and siWWC3-treated cells revealed cell elongation and moreover showed a reduction of jagged and reticular junctions and increased smoothed cell borders (**Fig. 5B1-3**). Of note, CDH5 intensity was relatively unchanged. Altered cell morphology was also observed staining for f-actin (phalloidin, or Phal), where cell elongation in siWWC3 cells was accompanied by enrichment of cortical actin and reduction of stress fibers (**Fig. 5B4-6**).

**Figure 5.**
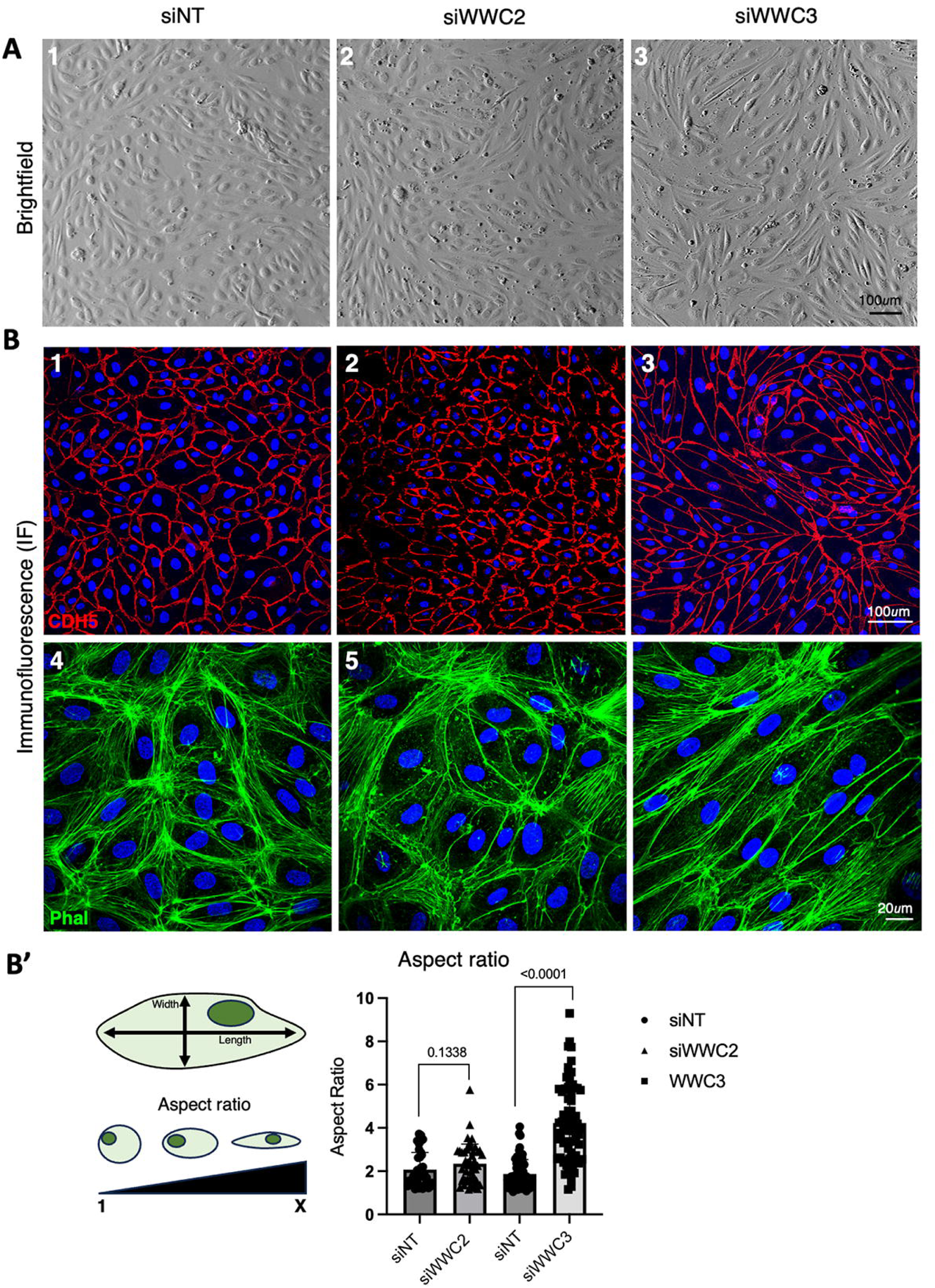
Loss of WWC3, but not WWC2, impacts EC cell shape and organization. **A)** Brightfield and **B)** CDH5 or Phallodin Immunofluorescence (as indicated) reveal EC elongation upon loss of WWC3 compared to siNT and siWWC2 treated ECs. Aspect ratio of cells in B is analyzed to quantify cell elongation is shown in **B’**. Significance was calculated using unpaired Student’s t-test. Scale bars: Panels A1-3 and B1-3 are 100μm, Panels B4-6 are 20μm.

### WWC3 is required to maintain endothelial identity

Increased nuclear YAP/TAZ has been linked to higher expression in TGFβ2, leading to endo-to-mesenchymal transition (EndoMT).^5,23,24^ Given that siWWC3 ECs displayed significantly altered cell shape, reminiscent of a fibroblastic, mesenchymal cell type, we investigated whether increased YAP1/TAZ signaling might lead to increased TGFβ signaling in ECs. Using qPCR to examine expression of classical EMT genes, including *ACTA2*, *FBN*, *CDH2,* and *VIM,* we assessed both siWWC2 and siWWC3-treated ECs. We observed that endothelial markers like *CDH5* and *vWF* did not change upon loss of *WWC2*, suggesting that endothelial fate did not change, and nor did mesenchymal gene expression (**Fig. 6A**). However, *vWF* expression decreased in siWWC3-treated cells, along with a concomitant increase in some mesenchymal markers, including fibronectin (*FBN*) and N-cadherin (*CDH2*). Interestingly, vimentin (*VIM*), a known mesenchymal transition driver, showed instead a significant reduction in expression levels in ECs upon *WWC3* depletion.

**Figure 6.**
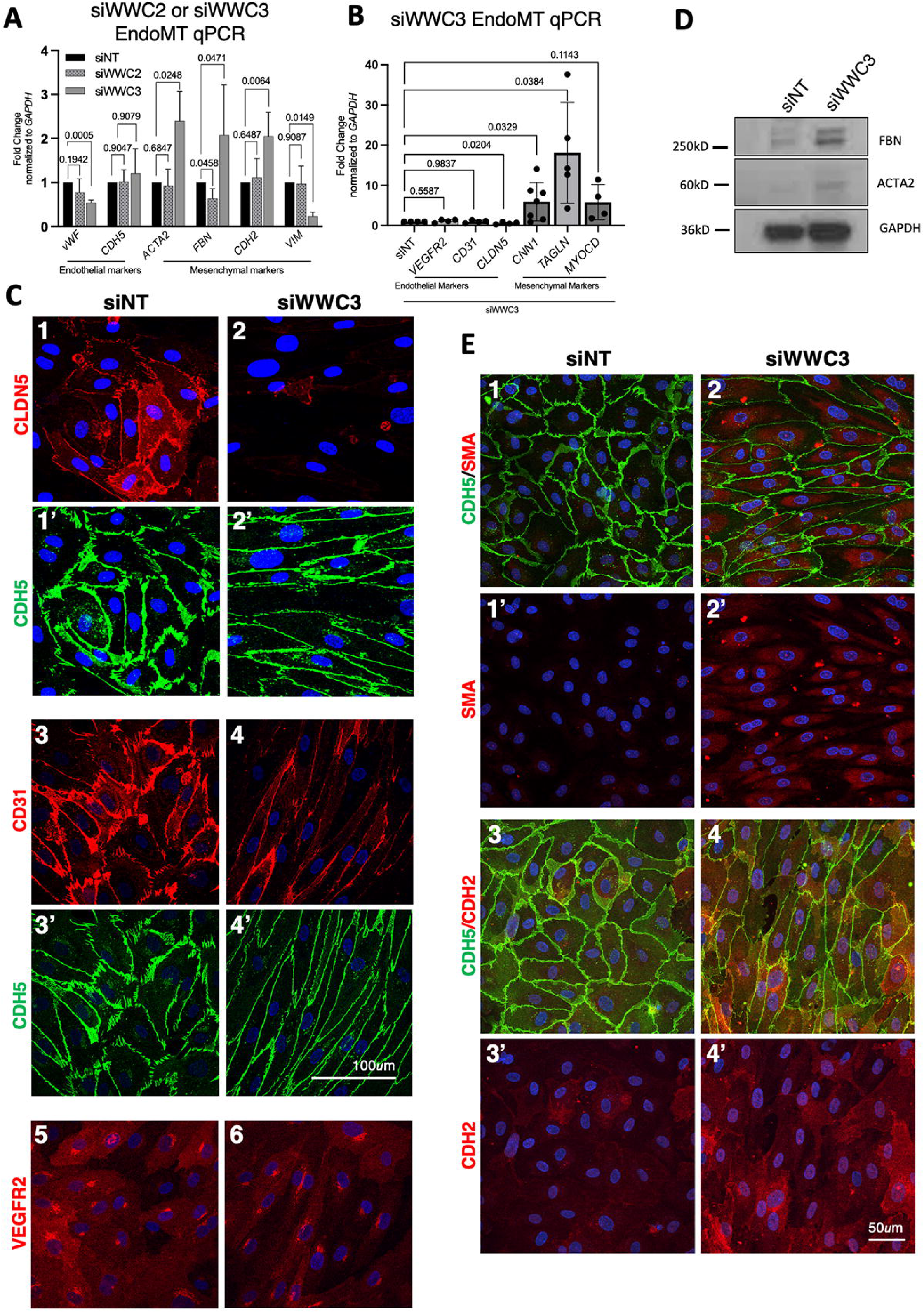
Loss of WWC3 upregulates EMT markers in ECs. **A)** qPCR data shows the increase in mesenchymal genes upon loss of WWC3 in ECs, except *VIM*, which is reduced, but not in siWWC2 ECs. **B)** Analysis by qPCR showing increase in additional mesenchymal genes in siWWC3. Endothelial genes by contrast are largely unchanged. **C)** Immunofluorescence staining shows that ECs junctional markers like CLDN5 and CD31 are reduced in *WWC3* knockdown. VEGFR2 expression by contrast is unchanged. **D)** Increase in protein levels of mesenchymal genes is shown by Western blot in siWWC3 treated ECs. **E)** Immunofluorescence staining shows the upregulation of mesenchymal markers like SMA and CDH2 in siWWC3 cells. Significance was calculated using unpaired Student’s t-test. Scale bars: Panels C1-6 are 100μm, Panels E1-4’ are 50μm.

We expanded our analysis of *WWC3-*depleted cells and further analyzed other endothelial and mesenchymal markers. We observed a striking decrease in *Cldn5* levels, though no change in *VEGFR2* or *CD31*, and a marked upregulation of *CNN1* and *TGLN* (**Fig. 6B**). Similarly, examination of endothelial markers by IF confirmed the partial loss of endothelial markers that we observed at the transcript level. We observed a dramatic reduction in the protein levels of the EC junction marker *CLDN5* and *CD31*, but not *VEGFR2* (**Fig. 6C**). Interestingly, loss of *CD31* from the junctions in siWWC3-treated cells is not due to change in its transcript levels. The partial mesenchymal state of siWWC3 cells was further corroborated using Western blot analysis that revealed an increase in FBN and ACTA2 (smooth muscle actin, or SMA) protein (**Fig. 6D**). This could also be observed by using IF, as both SMA and N-cadherin (CDH2), where increased staining was evident in the cytoplasm of siWWC3 cells (**Fig. 6E**). Taken together, our results suggest that WWC3 is required for ECs to stably maintain their endothelial fate and prevent their transitioning into a mesenchymal fate, while WWC2 is likely not required for this process.

## DISCUSSION

Although WWC family proteins are widely regarded as functionally redundant upstream activators of the Hippo pathway, we find that two WWC proteins in human ECs perform distinct functions. We show that WWC3 is the predominant regulator of canonical Hippo signaling in human ECs, exerting a greater influence on LATS1/2 phosphorylation and YAP1/TAZ activity. Of great interest, it also essential for maintenance of endothelial identity. By contrast, WWC2 has relatively modest effects on canonical Hippo signaling, but plays a more prominent role in regulating responses to VEGF. Despite these distinct molecular functions, depletion of WWC2 or WWC3 impaired fundamental endothelial behaviors, including proliferation, migration and cord formation, indicating that both proteins contribute to endothelial function via different mechanisms. Together, our findings challenge the prevailing view that WWC proteins function interchangeably and instead support a model whereby WWC family proteins have undergone functional specialization in the endothelium.

### Functional specialization of WWC2 and WWC3 in endothelial cells

The WWC family of proteins have been widely studied in the context of Hippo Signaling as facilitators of LATS1/2 phosphorylation.^14^ WWC1 is well-studied due to its role in cognitive function and association with Alzheimer’s Disease.^25,26^ Owing to the largely assumed redundancy between WWC proteins, WWC2 and WWC3 have received less attention. Thus, the distinct functions of WWC2 and WWC3 in human ECs are somewhat unexpected given the biochemical similarities among WWC family proteins. In HUVECs, however, depletion of WWC3 resulted in a substantially greater reduction in LATS1/2 phosphorylation and a more pronounced effect on YAP1/TAZ localization and signaling than depletion of WWC2. These findings suggest that the ability of WWC proteins to perform similar biochemical functions in several cell types does not necessarily translate into equivalent roles in ECs.

Functional divergence in the WWC proteins has previously been seen in synapse formation, where WWC1 and WWC2 are responsible for AMPA-receptor and GABA-receptor localization at the synapse, respectively.^27^ Our studies further highlight this divergence by showing the while WWC2 has a modest effect on proliferation, WWC3 loss drastically decreases proliferation. Although high YAP/TAZ activity is associated with higher proliferation, our results suggest that higher nuclear YAP/TAZ in siWWC3 treated cells pushes the cells towards a mesenchymal state instead. The basis for this functional specialization remains unclear but could reflect endogenous differences in protein levels, subcellular localization, or interactions with distinct signaling partners. WWC proteins function as molecular scaffolds, making their activity particularly dependent on the proteins and cellular compartments with which they associate. Hence, WWC2 and WWC3 may assemble distinct signaling complexes in ECs, allowing closely related proteins to regulate different aspects of endothelial cell behavior.

### WWC function in mouse and human ECs

The prominent role of WWC3 in human ECs is particularly interesting when placed in the context of previous genetic studies in mice. Loss of *Wwc2* in mice causes severe defects in embryonic vascular development,^15^ while post-natal EC-specific deletion of *Wwc2* results in milder aberrant angiogenic sprouting.^16^ By contrast, loss of Wwc1 alone in mouse does not result in overt vascular defects.^16^ These studies have thus pointed to WWC2 as the major WWC family member important in vascular development in mice. Our findings suggest the situation may differ in human ECs, where WWC3 is broadly expressed and has a substantially greater effect than WWC2 on LATS1/2 phosphorylation and downstream YAP1/TAZ signaling. Notably, WWC3 is absent from the mouse genome, raising the possibility that functions carried by WWC2 in mouse endothelium may be distributed between WWC2 and WWC3 in human. Whether this represents an evolutionary divergence and functional specialization of WWC family members or reflects differences in expression and cellular context is still unclear at this point. However, our findings highlight an interesting distinction between mouse and human endothelial Hippo regulation and suggest that the vascular functions of WWC2 described in mouse may not fully predict how WWC proteins function in human endothelium. Further studies are required to ascertain the role of WWC3 in human physiology to understand the regulation of Hippo signaling and YAP/TAZ activity in various pathologies.

### WWC regulation of VEGF signaling

The modest effect of WWC2 depletion on canonical Hippo signaling in human ECs raises the question of what function WWC2 has in this cell type. Our findings point to regulation of VEGF signaling as one potential role. Whereas VEGF-induced phosphorylation of VEGFR2 and its downstream effectors declined over time in control ECs following VEGF treatment, loss of WWC2 resulted in prolonged VEGFR2 and ERK1/2 activation, suggesting that WWC2 contributes to the normal attenuation of VEGF signaling. This finding is consistent with studies showing that the upstream Hippo regulator Merlin/NF2 controls angiogenic sprouting by regulating VEGFR2 internalization independently of YAP1/TAZ.^9^ WWC proteins are molecular scaffolds that are often membrane-associated and play roles in trafficking, raising the possibility that WWC2 similarly influences VEGFR2 internalization, recycling and degradation rather than primarily acting through the canonical LATS and YAP1/TAZ axis. Our observations, however, do not distinguish between these possibilities, and WWC3 depletion also resulted in persistence of VEGF signaling at later time points. This suggests that regulation of this pathway might not be exclusive to WWC2. Nevertheless, the more pronounced effect of WWC2 loss on the VEGF signaling response supports the idea that WWC family members have distinct signaling roles in ECs.

### Endothelial identity requires WWC3

In contrast to the effects of WWC2 on VEGF signaling, loss of WWC3 produced a significant change in EC morphology and fate. WWC3 depletion strongly reduced LATS1/2 phosphorylation and increased nuclear YAP1/TAZ along with an increase in TGFβ2 expression. These molecular changes coincided with elongation of ECs, altered cytoskeleton, loss of *CLDN5* and *vWF,* loss of *CD31* from EC junctions and increased expression of some mesenchymal markers, including *CNN1*, *TAGLN*, *FBN* and *ACTA2*. Together, our observations suggest that WWC3-dependent Hippo signaling contributes to maintenance of endothelial identity and that its loss promotes an EndoMT-like shift, with acquisition of mesenchymal characteristics. This is consistent with YAP1/TAZ and TGFβ signaling regulating cell fate. In addition, the strong induction of TGFβ upon WWC3 knockdown potentially links reduced LATS activity and the cellular changes observed. However, siWWC3 cells do not undergo complete EndoMT, as several endothelial key genes are still expressed, and only a subset of mesenchymal markers are aberrantly induced. We therefore propose that loss of WWC3 induces a partial EndoMT-like state, rather than a complete mesenchymal conversion. We note that classical EndoMT is associated with increased cell motility, while WWC3 loss in HUVECs decreases motility, demonstrating that siWWC3 cells have not fully shifted to a mesenchymal fate. Interestingly, the loss of WWC2 also results in the increase of TGFβ2 although to a much lesser extent, and yet it does not induce mesenchymal genes, the reason for this remains to be explored.

## Conclusion

Together, our study demonstrates that WWC2 and WWC3 perform overlapping but distinct functions in human ECs. Instead of acting like interchangeable upstream Hippo pathway scaffolds, WWC3 plays a predominant role in canonical LATS-YAP1/TAZ signaling and maintenance of EC identity. WWC2, on the other hand, has a comparatively greater impact on VEGF signaling dynamics. These observations reveal an unexpected functional specialization among WWC family members and underscore potential differences in regulation of endothelial Hippo signaling between mice and humans. More broadly, they also raise the possibility that WWC proteins integrate Hippo signaling with other pathways that control EC behavior, morphology and blood vessel homeostasis. As WWC2 is the only member of the family that is expressed in mouse vascular cells (data not shown), in vivo studies are required to understand the mechanisms by which it regulates angiogenesis. Given that WWC2 does not have a prominent effect on YAP/TAZ activity, at least in human ECs, it is important to identify mechanisms by which it regulates EC behavior in humans and how such mechanisms have been adapted in mouse during evolution.

## MATERIALS AND METHODS

### Cell culture and reagents

Primary human umbilical vein endothelial cells (HUVEC) between passages 3 to 7were grown in endothelial growth medium (EGM-2) containing 5% heat-inactivated fetal bovine serum (FBS) and endothelial cell growth supplements (Lonza). Cell cultures were maintained at 37°C in a 5% CO2 incubator with 95% humidity. siRNA transfection was carried out using Lipofectamine (Invitrogen). WWC2 and WWC3 were knocked down using Invitrogen SilencerSelect WWC2 siRNA (id: s36823; Catalogue: 4392420) at 50nM and Invitrogen Silencer WWC3 siRNA (id: 126853 Catalogue: AM16708) at 50nM respectively. Assays were carried out 48hrs to 72hrs post-transfection.

### Migration assay

Cells treated with siRNA were plated on Culture-Insert 3 Well in µ-Dish 35 mm (Ibidi: 80366) at 10,000 cells per insert in EGM2 complete media. Cells were starved for 4 hours in EBM2. Inserts were removed and EBM2 + 2% FBS was added. Cells were imaged at this time-point for initial wound area and then at 16 hours for the final wound area using Keyonce. Percentage wound closure was calculated from 3 experiments with 3 to 4 FOVs per wound accordingly.

### Tube formation assay

Cells treated with siRNA were plated on Matrigel (Corning: 356231) in 96-well plate with cover slip bottom (Cellvis: P96-1.5H-N) at 25000 per well in EGM2 complete media. After 12 hours, cells were stained with Calcein AM (Invitrogen: C3099) at 2ug/mL for 30 minutes in the incubator and then imaged. Each condition was plated in triplicate and the number of nodes were counted for 3 separate experiments.

### Immunofluorescence assays

HUVECs were fixed with 4% paraformaldehyde for 10 minutes at room temperature or cold methanol on ice for 2 minutes. 0.01% TritonX-100 (ThermoFisher: X100) in PBS was used for permeabilization for 10 minutes at room temperature and followed by 1 hour blocking with CAS-Block (Thermofisher: 008120). Cells were incubated in primary antibodies overnight and in secondary antibodies in CAS-Block for 1hr. Antibodies are indicated in the table below (**Table 1**).

### VEGF signaling assays

Cells treated with siRNA were plated on 6-well plates and after 24 hours, they were starved for 1 hour using EBM2 and then treated with VEGF at 50ng/mL (R&D 293-VE-010/CF) in EBM2 for indicated time points. Protein lysates were collected and used for Western Blot analysis.

### Protein extraction and Western blot

Cells were rinsed with ice-cold PBS, then lysed in RIPA buffer (Abcam) containing protease and phosphatase inhibitors (ThermoFisher catalog: A32961) according to manufacturer’s instructions. Protein concentration was determined using the Pierce BCA protein assay kit. Subsequent Western blot analysis was done as previously described (Cowdin et al., 2025). Antibodies are indicated in the table below.

### RNA extraction and quantitative PCR

Real-time qPCR analysis was done as previously described (Cowdin et al., 2025). Briefly, RNA was extracted using RNeasy Mini Kit (Qiagen) and cDNA was synthesized using Super-Script III (Invitrogen). qPCR was performed with SYBR Green Master Mix (Applied Biosystems) using gene-specific primers (**Table 2**).

### Quantification and statistical analysis

#### Proliferation assay

20 to 30 FOVs were used to calculate the percentage of Ki67-positive nuclei across three separate experiments per condition.

#### Cell morphology

Five cells from 10 FOVs from three separate experiments were traced using the CDH5 stain to measure their major and minor axis, and the cell elongation was measured by the aspect ratio.

#### YAP1/TAZ Nucleus to cytoplasmic ratio and Phospho-YAP1 Intensity

Mean gray value of the nucleus and the cytoplasm 100 cells from 10 to 16 FOVs across three separate experiments per condition were used for calculating the ratio and the cytoplasmic pYAP1 intensity.

Data were plotted and analyzed using GraphPad 10.2.3 Prism. Dot plots were plotted to accurately depict the spread of the data. Significance was calculated using unpaired Student’s t-test. A P<0.05 was considered significant. All bars are mean ± SEM from at least 3 different experiments unless stated otherwise. Figures and model were made using Microsoft PowerPoint.

## ACKNOWLEDGEMENTS

We are grateful to the members of the Cleaver lab for their thoughtful commentary and discussions in experimental interpretation and the preparation of this manuscript. We appreciate the assistance of the UTSW Quantitative Light Microscopy Core facility. In addition, we are grateful to Lu Sun (UTSW) and his lab for sharing expertise and equipment.

## Author Contribution

TP and OC conceptualized the study. TP and AM carried out cell culture experiments. TP performed cell imaging and analysis. OC provided resources. TP and OC curated data. TP generated figures and OC edited them. TP and OC wrote the original draft of the manuscript, which was reviewed and edited by AM. OC acquired funding.

## SUPPLEMENTARY MATERIALS

**Table 1.** List of Antibodies.

| **Primary Antibody** | **Catalog No.** | **Dilution** |
| --- | --- | --- |
| WWC2 Mouse Antibody | Sc-515892 | 1:1000 (WB) |
| WWC3 Rabbit Antibody | HPA039814 | 1:1000 (WB) |
| Phospho-LATS1/LATS2 (Thr1079, Thr1041) Rabbit Monoclonal Antibody | MA569915 | 1:1000 (WB) |
| LATS1 (C66B5) Rabbit Monoclonal Antibody | 3477s | 1:1000 (WB) |
| Abcam Rabbit LATS2 | ab243657 | 1:2000 (WB) |
| YAP (D8H1X) Rabbit Monoclonal Antibody | 14074 | 1:1000 (WB) |
| TAZ Rabbit Antibody | HPA039557 | 1:2000 (WB) |
| YAP/TAZ (D24E4) Rabbit Monoclonal Antibody | 8418 | 1:100(IF) |
| Phospho-YAP (Ser127) Antibody | 4911 | 1:1000 (WB) |
| Abcam Anti-Ki67 antibody [SP6] | 16667 | 1:400 (IF) |
| Human VE-Cadherin Goat Antibody | AF938 | 1:400 (IF) |
| Human PECAM Mouse Antibody | 555444 | 1:200 (IF) |
| Phospho-VEGFR2 (Tyr 1175) Rabbit Antibody | 2478 | 1:1000 (WB) |
| VEGFR2 Rabbit Antibody | 2479 | 1:2500 (WB)<br>1:100 (IF) |
| Phalloidin Alexa Fluor 488 | AF12379 | 1:800 (IF) |
| Phospho-AKT (Ser 473) Rabbit Antibody | 4058 | 1:1000 (WB) |
| AKT Rabbit Antibody | 9272 | 1:2500 (WB) |
| Phospho-ERK1/2 (Thr 202/ Tyr 204) Rabbit Antibody | 9101 | 1:5000 (WB) |
| ERK1/2 Rabbit Antibody | 4695 | 1:5000 (WB) |
| Smooth Muscle Actin Mouse Antibody | ab7817 | 1:2500 (WB) |
| Fibronectin Mouse Antibody | 610078 | 1:1000 (WB) |
| CDH2 Mouse Antibody | 690120 | 1:100 (IF) |
| Ki67 Rabbit Antibody | ab16667 | 1:400 (IF) |
| GAPDH Rabbit Antibody | 2118 | 1:10000 (WB) |
| HSP90 Rabbit Antibody | 4874 | 1:5000 (WB) |

| Secondary Antibody | Catalog No. | Dilution |
| --- | --- | --- |
| Anti-Rabbit HRP | HAF008 | 1:5000 (WB) |
| Anti-Mouse HRP | ab97030 | 1:5000 (WB) |
| Anti-Rabbit 488 | A-21206 | 1:400 (IF) |
| Anti-Rabbit 555 | A-31572 | 1:400 (IF) |
| Anti-Rabbit 647 | A-31573 | 1:400 (IF) |
| Anti-Mouse 488 | A-21202 | 1:400 (IF) |
| Anti-Mouse 555 | A-31570 | 1:400 (IF) |
| Anti-Goat 647 | A-21447 | 1:400 (IF) |
| Anti-Goat 555 | A-21432 | 1:400 (IF) |
| Anti-Goat 488 | A-11055 | 1:400 (IF) |

**Table 2.**
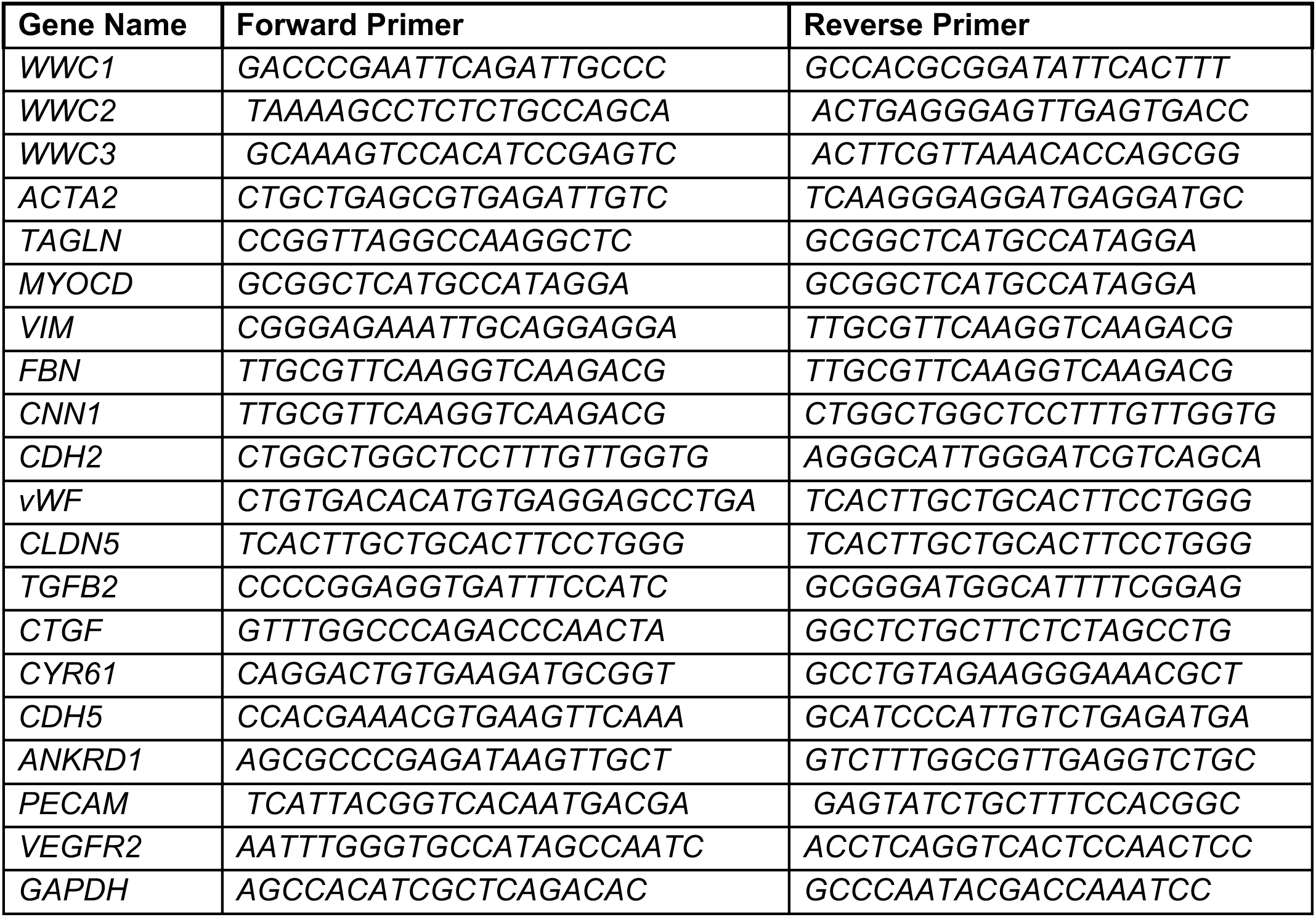
List of Primers.

